# HSV-1 exploits host heterochromatin for egress

**DOI:** 10.1101/2022.05.31.494218

**Authors:** Hannah C Lewis, Laurel E Kelnhofer-Millevolte, Mia R Brinkley, Hannah E Arbach, Edward A Arnold, Srinivas Ramachandran, Daphne C Avgousti

## Abstract

Herpes simplex virus (HSV-1) progeny form in the nucleus and must exit to successfully infect other cells. These newly formed viral capsids navigate the complex chromatin architecture of the nucleus to reach the inner nuclear membrane and egress. Here, we demonstrate by transmission electron microscopy (TEM) that HSV-1 capsids traverse dense heterochromatin in the nuclear periphery to reach the inner nuclear membrane. We found that this heterochromatin is dependent on the specific chromatin marks of trimethylation on histone H3 lysine 27 (H3K27me3) and the histone variant macroH2A1. Through chromatin profiling over the course of infection, we revealed massive host genomic regions bound by macroH2A1 and H3K27me3 that correlate with decreased host transcription in active compartments. This indicates the formation of new heterochromatin during infection. We found that loss of these markers resulted in significantly lower viral titers but did not impact viral genome or protein accumulation. Strikingly, we discovered by TEM that loss of macroH2A1 or H3K27me3 resulted in nuclear trapping of viral capsids. Thus, our work demonstrates that HSV-1 takes advantage of the dynamic nature of host heterochromatin formation during infection for efficient viral egress.

## INTRODUCTION

Nuclear replicating viruses must contend with host chromatin to establish a successful infection. Like most DNA viruses, herpes simplex virus (HSV-1) takes advantage of host chromatin factors both by incorporating histones onto its genome to promote gene expression^1^ and by reorganizing host chromatin during infection^2–4^. This redistribution of host chromatin results in a global increase in heterochromatin^5^. Host chromatin that accumulates in the nuclear periphery during HSV-1 infection creates a potential barrier for capsids to egress from the nucleus. HSV-1 capsids egress by a unique mechanism of budding into the inner nuclear membrane and then fusing with the outer nuclear membrane for further maturation in the cytosol^6,7^. Viral capsids traversing this dense chromatin in the nuclear periphery are associated with channels of less dense staining, termed heterochromatin channels^8^. However, it is not known if these channels are necessary for viral egress and the mechanisms by which they form are unclear.

Heterochromatin density and subnuclear localization is affected by the presence and ratio of specific histone modifications. In uninfected cells, histone modifications such as trimethylation of histone H3 lysine 27 (H3K27me3) and the histone variant macroH2A1, among others, delineate heterochromatin regions that are largely localized to the nuclear periphery. H3K27me3 is deposited by the EZH2 enzyme^9^, a member of the polycomb repressive complex (PRC2)^10^. This modification is bound by PRC2, leading to modification of adjacent nucleosomes, which results in formation of dense heterochromatin domains that repress transcription. MacroH2A, the largest of the histone variants, consists of a canonical histone fold domain, a small linker region and a C-terminal 25kDa macro domain that is thought to protrude from the nucleosome^11^. There are three isoforms referred to collectively as macroH2A: macroH2A1.1, macroH2A1.2 and macroH2A2. MacroH2A1.1 and macroH2A1.2 are splice variants of the same gene that differ by one exon resulting in a 28-amino acid difference in the macro domain. MacroH2A1 was found to be downregulated in melanoma^12^, suggesting a key role in the maintenance of genome integrity. Importantly, loss of macroH2A1 and macroH2A2 results in a significant decrease in heterochromatin in the nuclear periphery observed by electron microscopy^13^. Furthermore, macroH2A1 also demarcates regions of host chromatin that associate with the nuclear lamina^14^, termed lamina-associated domains (LADs), highlighting its importance in linking chromatin with the nuclear envelope to support nuclear integrity.

In this study, we visualized HSV-1 nuclear egress by transmission electron microscopy (TEM) and found that capsids reach the inner nuclear membrane in regions of less densely stained chromatin. Therefore, we hypothesized that HSV-1 exploits host heterochromatin to generate channels and successfully egress from the nuclear compartment. We examined chromatin structure by TEM in the absence of heterochromatin markers macroH2A1 and H3K27me3 and discovered that peripheral heterochromatin is largely dependent on these marks. We used chromatin profiling of macroH2A1 and H3K27me3 during HSV-1 infection to define the specific host genomic regions bound by these markers and found that they demarcate broad regions of heterochromatin that form in transcriptionally active compartments. Importantly, we found that the loss of macroH2A1 results in significantly lower viral titers but does not impair viral transcription, protein production, or replication in both lab adapted and clinical isolates of HSV-1. Furthermore, by inhibiting EZH2 deposition of H3K27me3, we found that reduction of H3K27me3 also leads to a significant decrease in viral titers. Finally, we determined by TEM that loss of macroH2A1 or H3K27me3 results in significantly more viral capsids trapped in the nuclear compartment, pinpointing the importance of heterochromatin in viral nuclear egress. Our study is the first to demonstrate that HSV-1 infection manipulates heterochromatin markers to successfully egress from the nuclear compartment.

## RESULTS

### HSV-1 capsids associate with regions of less dense chromatin to escape the nucleus

To investigate the journey of HSV-1 capsids to the inner nuclear membrane, we used transmission electron microscopy (TEM) to image nuclei and examined heterochromatin formation in human foreskin fibroblast (HFF) cells. HFF cells are used as a model for HSV-1 infection as they represent a common site of infection in humans^15^. In uninfected cells, we observed the characteristic dark heterochromatin staining in the nuclear periphery (Fig 1a, arrowhead). Upon infection with HSV-1, we observed capsids interacting with the inner nuclear membrane primarily in regions of less dense staining (Fig 1b, arrows). These results are consistent with previous reports in African green monkey kidney cells (Vero) and human B cells where viral capsids were observed to interact with the inner nuclear membrane in regions of less dense staining, or heterochromatin channels^8,16^. Because heterochromatin in the nuclear periphery was dependent on macroH2A presence in hepatoma cells^13^, and macroH2A1 commonly overlaps with H3K27me3^17^, we chose to examine heterochromatin formation in the absence of these markers in HFFs.

**Figure 1.**
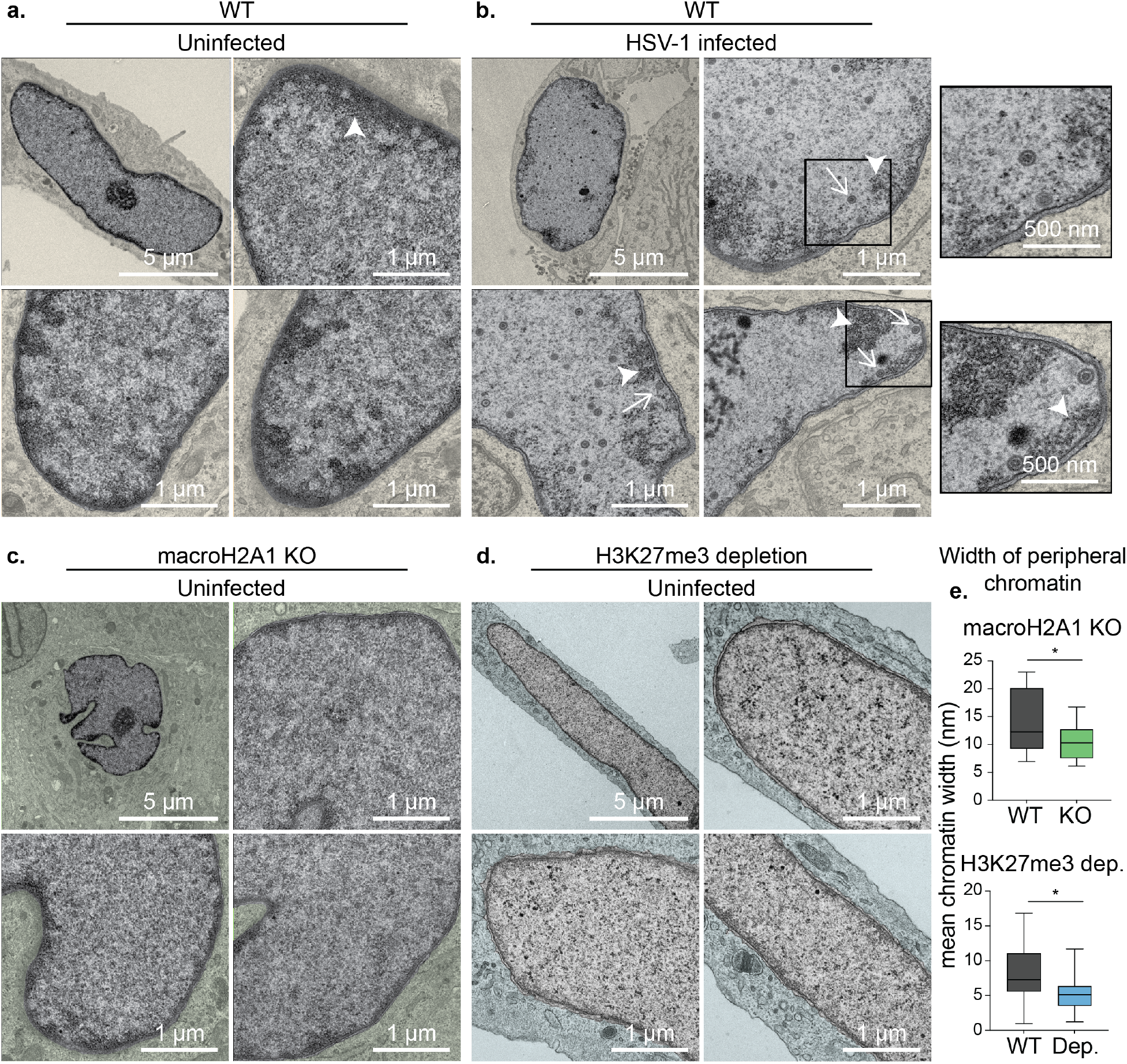
HSV-1 capsids traverse channels of less dense chromatin to reach the inner nuclear membrane. a) Transmission electron microscopy (TEM) images of representative uninfected nuclei in WT HFF-Ts. Regions outside of the nucleus are colorized yellow. Dark regions represent high density heterochromatin (arrowhead). Scale bar as indicated. b) TEM images of representative WT nuclei at 18 hours post-infection with HSV-1. Insets show enlarged views of respective boxed areas containing heterochromatin channels within nuclei. Arrowhead indicates heterochromatin, arrows indicate HSV-1 capsids. Scale bar as indicated. c) TEM images of representative uninfected nuclei in macroH2A1 knockout HFF-Ts. Regions outside of the nucleus are colorized green. Scale bar as indicated. d) TEM images of representative uninfected nuclei in H3K27me3 depleted HFF-Ts. Regions outside of the nucleus are colorized blue. Scale bar as indicated. e) (Top) Quantification of peripheral heterochromatin width in nuclei from A and C. Width was measured in nm from the nuclear periphery via binary thresholding from intensity profiles sampled every 10 pixels. Mean width was plotted for each nucleus. p= 0.0275 (n=16 WT, n=22 macroH2A1 KO). (Bottom) Quantification as above in nuclei from A and D (p= 0.0144, n= 22 WT, n=20 H3K27me3 depleted).

### Heterochromatin markers macroH2A1 and H3K27me3 are critical for heterochromatin formation in the nuclear periphery

We used CRISPR-Cas9 to knock-out (KO) expression of the *macroH2A1* gene in hTERT-immortalized HFF cells (HFF-T) cells, which resulted in the loss of macroH2A1.1 and macroH2A1.2, termed macroH2A1 KO cells. We found by TEM that macroH2A1 KO cells had strikingly less heterochromatin in the nuclear periphery than WT cells (Fig 1c). We quantified the width of heterochromatin at the nuclear periphery in each cell type through binary thresholding of the intensity of electron-dense regions and found that loss of macroH2A1 results in a significant reduction in peripheral heterochromatin (Fig 1e). To reduce H3K27me3 levels, we targeted the EZH2 enzyme that deposits this mark^18^ using a well-characterized inhibitor called tazemetostat (EPZ-6438)^19^. We treated cells with 10 µM of tazemetostat or DMSO control for 3 days to allow for reduction of H3K27me3 and imaged nuclei by TEM (Fig 1d). Similar to macroH2A1 KO cells, we found a significant decrease in heterochromatin in the nuclear periphery of cells with depleted H3K27me3 (Fig 1e). It is important to note that in cells with observed decreased heterochromatin that there is not a decrease in total chromatin. These cells are simply not able to condense their chromatin to a degree that it would be visible by TEM. Importantly, the genomic regions bound by macroH2A1 and H3K27me3 are sufficiently massive that changes are visible by TEM. Therefore, we next examined the genomic distribution of these markers by chromatin profiling.

### MacroH2A1 and H3K27me3 bind broad regions of the host genome that increase during infection

We hypothesized that macroH2A1 and H3K27me3 are deposited at specific genomic loci on the host genome during infection to promote the formation of heterochromatin. To test this hypothesis, we used CUT&Tag^20^ to profile the genomic localization of macroH2A1 at 4, 8 and 12 hours post infection (hpi) in wild-type (WT) and macroH2A1 KO HFF-T cells. We also examined the chromatin profile of H3K27me3 under these conditions. On the human genome, we observed clear enrichment of macroH2A1 and H3K27me3 compared to IgG in WT cells (Fig 2a and S1a-b). The enrichment of macroH2A1 and H3K27me3 was observed as large domains that were gained upon viral infection (Fig 2a), suggesting that the host landscape is altered upon infection. These gains were reflected in an increase in total protein levels measured by western blot (Fig 2b). Since large regions are not amenable to traditional peak-based analysis, we instead used domain-based analysis. With the minimum domain size of 1 kb, we observed ∼50,000 H3K27me3 domains and ∼70,000 macroH2A1 domains genome wide. We observed less than 10 macroH2A1 domains in the datasets generated from macroH2A1 KO cells, indicating that our algorithm was identifying robust domains (Fig S2a). We then calculated the change in enrichments of macroH2A1 domains and used k-means clustering during the time course of infection compared to mock to identify patterns of macroH2A1 gain or loss during infection (Fig 2c and S2b). With k=6, we observed two clusters to substantially gain macroH2A1 over the course of infection (Clusters 5 and 6, Fig 2c left and S2d). Clusters 1-3 had significant decreases in macroH2A1, whereas cluster 4 had a minor increase (Fig 2c left, Figure S2d). We then asked how these macroH2A1-defined clusters behaved with respect to H3K27me3 and found that the overall trends across clusters were preserved in H3K27me3 (Fig 2c right and S2e), suggesting that H3K27me3 is largely enriched in the same broad regions. In contrast, the macroH2A1 CUT&Tag on the viral genome showed similar enrichments in both WT and macroH2A1 KO cells. Further, the H3K27me3 signal on the viral genomes also mirrored IgG control. These results indicated that there was a significant background signal from the viral genome that could not be accounted for (Fig S1c-d). Therefore, we disregarded viral genome reads in our dataset. Taken together, these results indicate that multiple forms of heterochromatin were gained at specific host genomic loci during HSV-1 infection.

**Figure 2.**
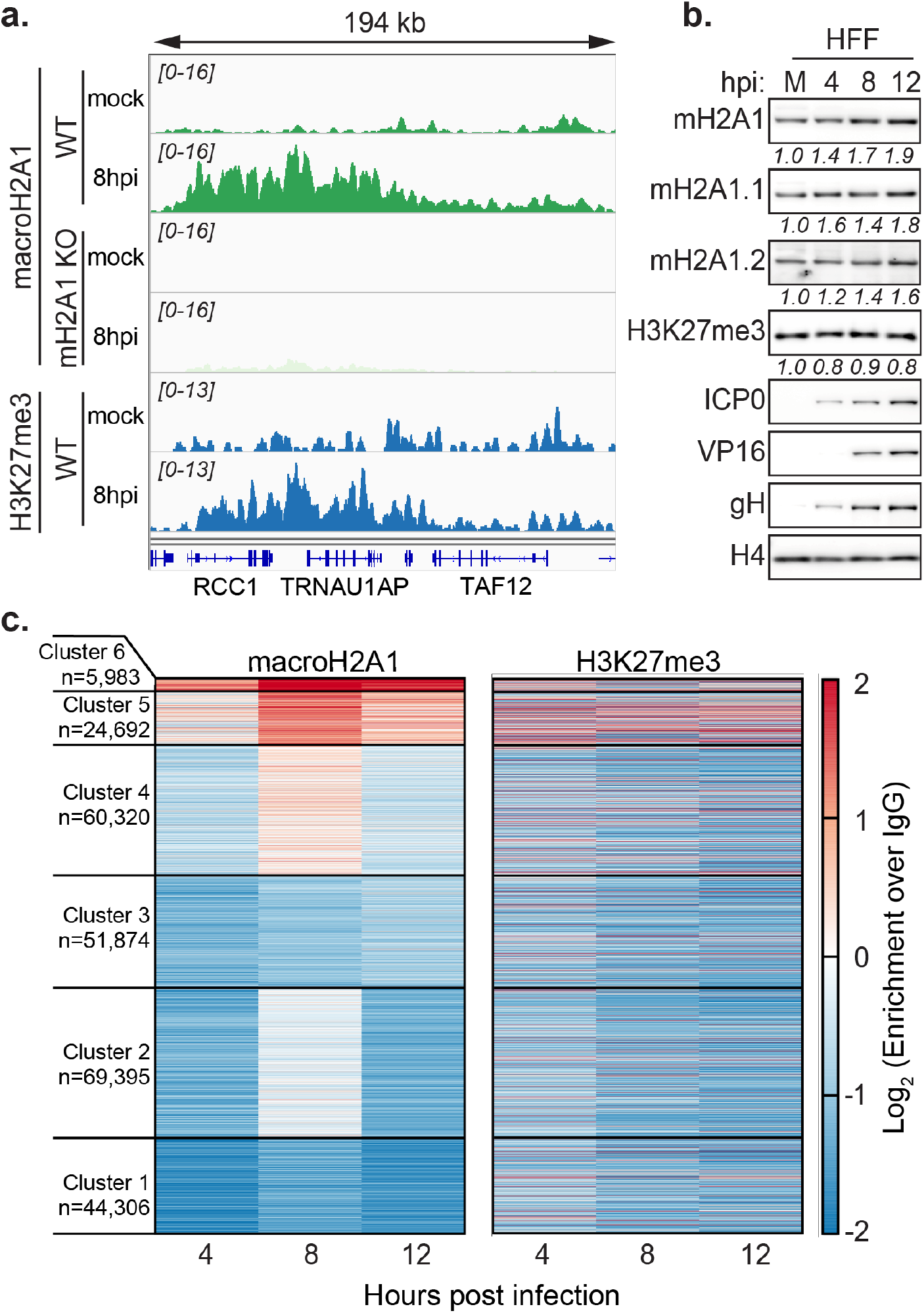
MacroH2A1 and H3K27me3 bind broad chromatin regions on the host genome. a) Representative genome browser snapshots of spike-in normalized CUT&Tag enrichment of macroH2A1 and H3K27me3 showing increases at 8hpi of HSV-1 infection as measured by CUT&Tag in WT or macroH2A1 KO as indicated. The region shown is found on chromosome 1. b) Western blot of total macroH2A1, macroH2A1.1, macroH2A1.2 isoforms, and H3K27me3 during mock (M) or HSV-1 infection 4-, 8-, and 12-hpi. ICP0, VP16, gH are viral proteins, H4 is shown as a loading control. Quantification of relative intensity of H4 normalized to the mock samples are shown. c) Changes in log2 enrichment of spike-in normalized CUT&Tag of macroH2A1 and H3K27me3 over IgG compared to mock treatment are shown as a heatmap. Each line in the heatmap represents a domain of macroH2A1.

To determine whether H3K27me3 deposition was dependent on macroH2A1, we also examined H3K27me3 domains in the absence of macroH2A1. We plotted the H3K27me3 changes at the same clusters for macroH2A1 KO cells and found that the trends of H3K27me3 were similar between WT and macroH2A1 cells (Fig S2b, c and f). This result suggests that H3K27me3 deposition is independent of macroH2A1. In summary, HSV-1 infection results in formation of new heterochromatin domains that span ∼10-100s of kilobases. Importantly, our results demonstrate that heterochromatin accumulating in the nuclear periphery during HSV-1 infection represents *new* regions of heterochromatin that are macroH2A1 and H3K27me3 dependent.

### MacroH2A1 and H3K27me3 deposition correlates with decreased transcription in active compartments

To examine the transcriptional output of newly formed macroH2A1 and H3K27me3-bound regions, we performed RNA-seq on WT and macroH2A1 KO cells over the course of HSV-1 infection and calculated fold changes in RNA levels at each time point over the mock control. We identified genes that are contained within domains in each cluster defined in Figure 2 and plotted the distribution of RNA fold changes of the genes grouped by the clusters to which they belonged (Fig S3a). We found that total RNA levels anti-correlate with macroH2A1 presence: clusters 1-3 had an increase in RNA levels, whereas clusters 4-6 had a decrease in RNA levels over the course of infection (Fig S3a). Strikingly, the gene expression changes in macroH2A1 KO cells mirror that of WT cells, leading us to conclude that macroH2A1 deposition is not driving changes in total RNA (Fig S3b). To determine whether these changes in RNA reflected changes in transcription, we analyzed published 4sU-RNA-labeling data during HSV-1 infection^21^. We found that overall clusters 4-6 had increased signals compared to clusters 1-3, suggesting that active transcription is occurring in clusters 4-6. Importantly, the time course comparison revealed that in clusters 5 and 6, where we found an increase in macroH2A1 and H3K27me3 presence during infection, the 4sU-labeled RNA decreased at 8hpi compared to 4 hpi, indicating that the new heterochromatin is deposited in regions with active transcription (Fig S3c).

Interestingly, cluster 4 diverged between total RNA and 4sU-RNA data: 4sU-RNA increased whereas total RNA decreased. This indication of active transcription explains why cluster 4 features a weak macroH2A1 gain. Taken together, these results indicate that macroH2A1 and H3K27me3 presence correlates with a decrease in transcription in active regions.

Next, we asked if the gain or loss of macroH2A1 happened across previously defined genome compartments^22^. We determined the distribution of the eigenvector corresponding to A/B compartments for IMR90 Hi-C data from the 4DN project^23^. Here, positive values correspond to the A compartment, which features higher gene density and accessible or active chromatin, whereas negative values correspond to the B compartment, which features inactive chromatin. All clusters except cluster 2 are significantly biased towards one of the compartments: clusters 1 and 3 are significantly biased towards compartment B, whereas clusters 4-6 are significantly biased towards compartment A (Fig S3d). Strikingly, the median of the clusters increases moving from Cluster 1 to Cluster 6, correlating with macroH2A1 gain. Thus, macroH2A1 gains and losses happen in distinct genomic compartments upon HSV-1 infection. In summary, macroH2A1 and H3K27me3 presence correlates with decreased transcription over the course of HSV-1 infection in active compartments.

Because macroH2A1 and H3K27me3-bound regions span a large portion of the host genome, we asked whether any specific gene ontology (GO) categories were over-represented for the genes in each macroH2A1-defined cluster (Fig S4). Genes in clusters 1 and 2 (with decreased macroH2A1 and increased expression) were associated with response to dsRNA and inflammatory responses, as expected during viral infection. Surprisingly, clusters 4-6, with increasing macroH2A1, consisted mostly of housekeeping genes. This suggests that the deposition of macroH2A1 is not directly affecting the expression of specific genes that would be pro- or anti-viral, but rather it is likely that these changes in transcription are a result of the stress response to infection. Furthermore, we quantified total viral RNA via RNA-seq and determined that there were no significant changes in viral RNA accumulation between WT and macroH2A1 KO cells (Fig S3e), suggesting that viral transcription is not affected by the loss of macroH2A1. Therefore, given that heterochromatin in the nuclear periphery surrounds capsids that access the inner nuclear membrane (Fig 1), we considered the scenario in which the chromatin structure generated by newly formed macroH2A1 and H3K27me3 bound regions of the genome was important for viral egress. Therefore, we next examined infection progression in the absence of these markers.

### Loss of macroH2A1 or H3K27me3 results in reduced viral progeny but does not affect viral genome or protein accumulation

To determine the impact of macroH2A1 in HSV-1 infection, we infected WT and macroH2A1 KO HFF-T cells with HSV-1. Upon infection, we examined viral protein accumulation and observed similar levels of ICP0 (immediate early ^24^), VP16 (early ^25^) and glycoprotein H (gH, late ^26^) in both wild type (WT) and macroH2A1 KO cells (Fig S5a). We next examined HSV-1 replication and measured viral genome accumulation by droplet digital PCR (ddPCR). We observed no significant difference in viral genome accumulation between WT and macroH2A1 KO cells (Fig 3a, black and green bars), indicating that neither HSV-1 replication nor protein production is affected by macroH2A1 loss. These results are consistent with no significant changes to viral RNA as measured by RNA-seq (Fig S3e). To determine the impact of macroH2A1 loss on viral progeny production, we measured viral genome accumulation in the supernatant. We discovered a 9-fold decrease in the levels of HSV-1 genomes produced in the supernatant of macroH2A1 KO cells compared to those produced from WT cells at 12 hpi (Fig 3b, green and black bars). We further discovered that infectious progeny produced from macroH2A1 KO cells were significantly decreased compared to those produced from WT cells at 8 and 12 hpi (Fig 3c, green and black bars). This effect was consistent across cell types with similar results observed in macroH2A1 KO RPE cells (Fig S5b-c). Taken together, these data indicate that loss of macroH2A1 leads to a significant defect in productive viral progeny but not viral protein, RNA, or genome accumulation. Therefore, we conclude that macroH2A1 is important for HSV-1 egress.

**Figure 3.**
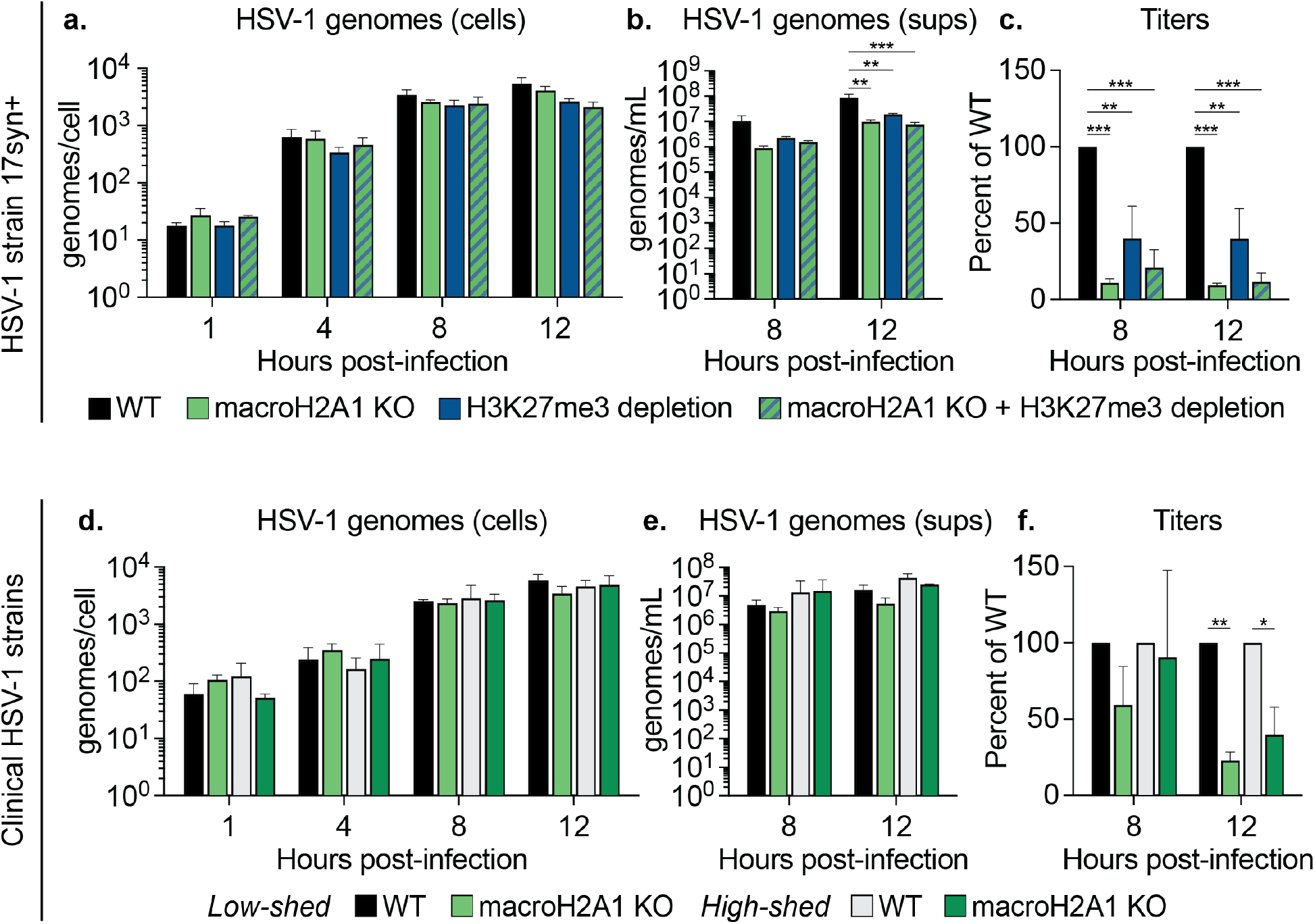
HSV-1 requires heterochromatin marks macroH2A1 and H3K27me3 for progeny production but not replication. a) Droplet digital (ddPCR) quantification of HSV-1 genomes extracted from infected cells as indicated for each time point. Error bars represent the SEM of three biological replicates. b) ddPCR quantification of HSV-1 genomes released from cells treated as indicated and isolated from supernatants (sups). Error bars represent the SEM of three biological replicates, ** p < 0.01, *** p < 0.001. c) Infectious progeny produced from HSV-1 infected cells treated as indicated and quantified by plaque assay. Viral yield is indicated as the percent yield compared to WT and was calculated through dividing plaque-forming units (pfu) per mL of the treatment group titers by the WT titers of each respective virus at each indicated time point. ** p < 0.01, *** p < 0.001. d) ddPCR quantification as in (a) from infection with low- and high-shedding clinical isolates of HSV-1. Error bars represent the SEM of three biological replicates. e) ddPCR quantification as in (b) of HSV-1 genomes from cells infected with clinical isolates of HSV-1 as indicated. Error bars represent the SEM of three biological replicates. f) Infectious progeny of cells as indicated infected with clinical HSV-1 isolates quantified by plaque assay. Viral yield is indicated as the percent yield compared to WT, * p. < 0.05, ** p < 0.01.

We next examined the impact of H3K27me3 on HSV-1 infection by treatment with tazemetostat to inhibit EZH2 and deplete H3K27me3 levels (as in Fig 1). We treated both WT and macroH2A1 KO cells to examine any synergistic effects of both heterochromatin markers. We found that accumulation of viral proteins ICP0, VP16 and gH were not affected by the reduction of H3K27me3, with or without macroH2A1 (Fig S5d-e). We next examined viral genome accumulation and found that reduction of H3K27me3 did not significantly affect viral genome accumulation compared to control (Fig 3a, black and blue bars). We observed no additional decrease in viral genomes in macroH2A1 KO cells treated with the inhibitor, suggesting that regardless of H3K27me3 levels, loss of macroH2A1 does not impact viral replication as we observed above (Fig 3a, striped bars). We next measured viral genomes in the supernatant to determine the impact of H3K27me3 reduction on progeny production and found a significant 4-fold decrease at 12 hpi from inhibitor-treated cells compared to DMSO-treated (Fig 3b, blue bars). Further, we found infectious progeny produced from inhibitor-treated cells were reduced significantly at 12 hpi compared to those produced from control cells (Fig 3c, blue bars). Interestingly, H3K27me3 reduction in macroH2A1 KO cells does not result in any additional defects, suggesting the two markers function in the same genetic pathway (Fig 3a-c, striped bars). These results are consistent with a previous report that found HSV-1 titers, but not viral replication, were reduced upon EZH2 inhibition^27^. Furthermore, we did not observe any significant differences in the ratio of genomes to plaque forming units (pfu) upon loss of macroH2A1, H3K27me3, or both, compared to control conditions (Fig S5f), indicating that loss of these markers does not result in significantly more defective particles. Taken together, these results indicate that H3K27me3 and macroH2A1 are important for efficient viral egress.

To determine how robust the requirement for macroH2A1-dependent chromatin is for HSV-1 egress, we used clinical HSV-1 viruses isolated from “low-shedding” or “high-shedding” patients. We measured protein accumulation at 4, 8, and 12 hpi and viral genome accumulation in cells and supernatant by ddPCR. We found that there was no significant change in viral protein or viral genome accumulation for either clinical isolate in macroH2A1 KO cells (Fig 3d and S5g-h). We measured viral genomes in the supernatant and found that there was a modest reduction in HSV-1 genomes in the supernatant of macroH2A1 KO cells compared to control cells (Fig 3e, green bars). For both clinical isolates, we found that macroH2A1 KO cells produced significantly fewer progeny than WT cells at 12 hpi (Fig 3f). Taken together, these results indicate that egress of clinically isolated HSV-1 is also dependent on macroH2A1.

### MacroH2A1 or H3K27me3-dependent heterochromatin in the nuclear periphery is critical for efficient HSV-1 nuclear egress

Our results indicate that HSV-1 capsids in the nucleus access the nuclear membrane through channels bracketed by highly dense chromatin at the nuclear periphery (Fig 1), consistent with previous reports in other cell types^8,16^. We found that loss of macroH2A1 or depletion of H3K27me3 resulted in decreased heterochromatin in the nuclear periphery (Fig 1) and caused a decrease in productive viral progeny (Fig 3). Therefore, we hypothesized that efficient HSV-1 egress out of the nuclear compartment requires macroH2A1- or H3K27me3-dependent heterochromatin in the nuclear periphery. To test this, we infected macroH2A1 KO cells, visualized nuclei by TEM and quantified capsids in the nucleus (Fig 4a). We found that infected macroH2A1 KO nuclei have significantly more HSV-1 capsids than those of WT cells (Fig 4a), indicating that loss of macroH2A1-dependent heterochromatin is detrimental for efficient nuclear egress. Similarly, we used tazemetostat treatment to deplete H3K27me3 levels, infected cells with HSV-1, and visualized capsids in the nuclei of infected cells by TEM. Strikingly, we found that significantly more capsids accumulated at the nuclear membrane compared to control cells (Fig 4b). We quantified this difference by scoring whether more than three capsids interacted with the inner nuclear membrane at any one location. Taken together, these results indicate that both macroH2A1 and H3K27me3-dependent heterochromatin in the nuclear periphery are crucial for successful viral nuclear egress.

**Figure 4.**
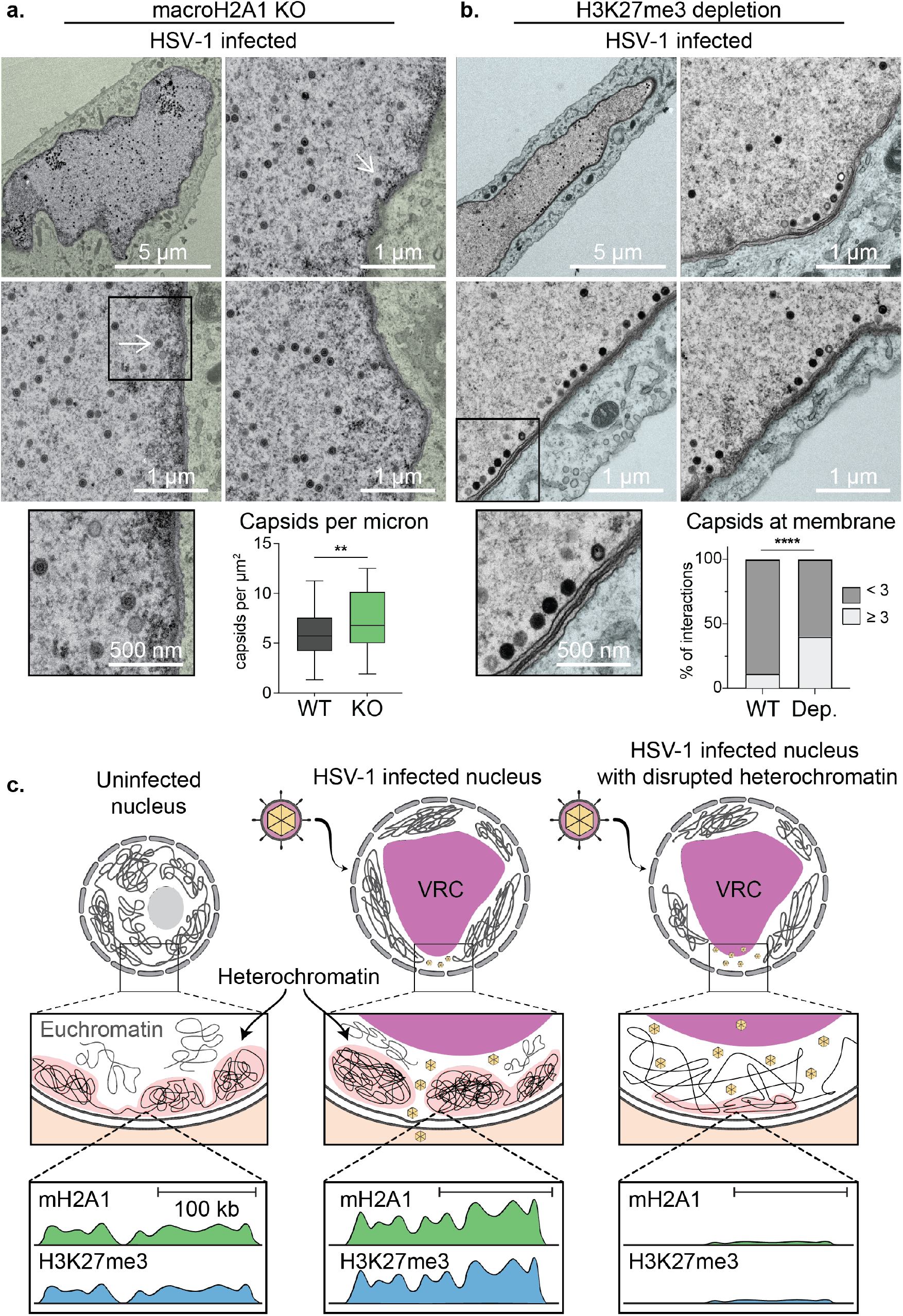
HSV-1 requires macroH2A1- and H3K27me3-dependent heterochromatin for efficient egress. a) TEM images of representative macroH2A1 KO nuclei infected with HSV-1. Insets show enlarged views of respective boxes. Arrows indicate HSV-1 capsids. Scale bar as indicated. (Bottom Right) Quantification of capsids within nuclei compared to Figure 1b. Number of capsids was normalized according to nucleus area in mm^2^, p=0.0036 (n=40 WT, n=55 mH2A1 KO). b) TEM images of representative H3K27me3 depleted nuclei infected with HSV-1 presented as in (a). (Bottom Right) Quantification of the number of capsids at the nuclear periphery compared to Figure 1b, p=0.0001 (n=8 cells for DMSO, n=5 cells for H3K27me3 depleted). c) Model for heterochromatin support of HSV-1 egress. In uninfected cells (left), heterochromatin is enriched in the nuclear periphery. Upon infection with HSV-1 (middle), there is an increase in heterochromatin at specific regions in the nuclear periphery that results in formation of heterochromatin channels that allow capsids to traverse from the viral replication centers (VRCs) to the inner nuclear membrane where they bud into the perinuclear space and subsequently fuse with the outer nuclear membrane into the cytoplasm. In infected cells missing heterochromatin markers macroH2A1 or H3K27me3 (right), these heterochromatin channels cannot form properly which results in increased capsids trapped in the nuclear compartment.

To determine whether capsid assembly was impacted by heterochromatin disruption, we quantified capsid type as a proportion of the total capsids from our TEM. Herpesvirus infection produces three subtypes of capsids: A capsids are empty, B capsids contain scaffolding proteins, and C capsids contain viral gDNA and are considered the precursor to infectious virions^28,29^ (Fig S6a). We quantified the proportion of each capsid type in WT and macroH2A1 KO cells and found no difference in the proportion of each capsid type (Fig S6b-c). We carried out the same analysis upon infection in H3K27me3-depleted cells and similarly found no difference in the proportion of capsid type (Fig S6c). In summary, these results indicate that capsid formation is not impacted by disruption of heterochromatin. Rather, the ability of the capsids to egress from the nuclear compartment is dependent on heterochromatin formation. Taken together, these data demonstrate that macroH2A1 and H3K27me3-dependent heterochromatin in the nuclear periphery is critical for efficient nuclear egress.

## DISCUSSION

Host chromatin is emerging as a critical barrier to viral success. Elegant electron microscopy work showed that HSV-1 infection induces host chromatin redistribution to the nuclear periphery^2,8^. Here, we showed that viral capsids subsequently egress through regions of less dense staining or heterochromatin channels (Fig 1), consistent with reports in different cell types^16^. We investigated the impact of heterochromatin markers histone variant macroH2A1 and H3K27me3 on chromatin architecture and HSV-1 infection. We found that loss of macroH2A1 resulted in significantly less heterochromatin visible in the nuclear periphery by TEM in HFF cells (Fig 1), similar to what was observed in hepatoma cells^13^. To our knowledge, this is the first instance of TEM imaging of heterochromatin upon depletion of H3K27me3, which produced a similar phenotype to macroH2A1 KO with significantly less heterochromatin in the nuclear periphery (Fig 1). We then demonstrated that macroH2A1 and H3K27me3 bind broad regions of the host genome that increase during HSV-1 infection (Fig 2). Our chromatin profiling analysis required a custom algorithm to accommodate the massive domains created by macroH2A1 and H3K27me3, consistent with regions large enough to be visible by TEM. We demonstrated that these regions are primarily found in transcriptionally active compartments defined by Hi-C, supporting the idea that newly formed heterochromatin increases during HSV-1 infection. We further showed that loss of macroH2A1 or depletion of H3K27me3 did not impact viral replication or protein production but resulted in significantly reduced infectious progeny (Fig 3). Finally, we demonstrated by TEM that in the absence of macroH2A1, significantly more viral capsids were trapped in the nucleus (Fig 4). Interestingly, the phenotype by TEM of H3K27me3 depletion on HSV-1 capsid accumulation was more striking with capsids lined up at the inner nuclear membrane (Fig 4). We found significantly more instances of greater than 3 capsids interacting with the nuclear membrane than in control cells, suggesting that capsids were unable to egress without H3K27me3-dependent heterochromatin. Together, these results support a model in which host heterochromatin formation acts as a structural highway for HSV-1 capsids to successfully access the inner nuclear membrane (Fig 4c).

Mining of proteomics on histone marks indicates that several heterochromatin marks including H3K9me3 are also changing during HSV-1 infection^3^. This suggests that there may be other mechanisms of heterochromatin dynamics at play. It will be fascinating to consider the diverse implications of heterochromatin domains not only as a means of transcriptional regulation but also as a structural component that impacts many facets of viral infection. Interestingly, a recent study examined chromatin accessibility changes upon HSV-1 infection compared to heat shock or salt stress and found that changes to chromatin were similar amongst these different conditions^30^. Together with our findings, this suggests that the global changes to heterochromatin are likely a consequence of the cell’s stress response to infection that HSV-1 exploits to access the inner nuclear membrane. Our results strongly indicate that it is the structural aspect of peripheral heterochromatin that is important for nuclear HSV-1 egress rather than transcription of any single gene or group of genes. Therefore, we propose that deposition of macroH2A1 and H3K27me3 in specific genomic regions during HSV-1 infection supports the formation of heterochromatin channels through which HSV-1 capsids travel to reach the inner nuclear membrane. At the inner nuclear membrane, capsids are aided by the nuclear egress complex, comprised of viral proteins UL34 and UL31, to bud into the inner nuclear membrane and fuse with the outer nuclear membrane for further maturation in the cytoplasm^7^. Budding into the nuclear membrane also requires nuclear lamina disruption. Multiple reports describe that not only does lamina disruption occur independently of capsid formation, but that chromatin is likely the limiting factor in viral egress^31–36^. Together with our results, this suggests that there are multiple steps in the capsid’s journey to the inner nuclear membrane preceding nuclear budding. In summary, our study demonstrates that the contribution of heterochromatin to the successful egress of HSV-1 capsids is a critical aspect of viral infection.

## AUTHOR CONTRIBUTIONS

Conceptualization, H.C.L, L.K.M., and D.C.A; methodology H.C.L, L.K.M., and D.C.A.; formal analysis, H.C.L, L.K.M., S.R., and D.C.A.; investigation, H.C.L, L.K.M., M.R.B., H.E.A, E.E.A, and D.C.A; resources, H.C.L, L.K.M., M.R.B., H.E.A, E.E.A, S.R., and D.C.A.; data curation, H.C.L, L.K.M., M.R.B., S.R., and D.C.A.; writing – original draft, H.C.L, L.K.M., M.R.B., S.R., and D.C.A.; writing – review & editing, H.C.L, L.K.M., M.R.B., H.E.A, E.E.A, S.R., and D.C.A.; visualization, H.C.L, L.K.M., S.R., and D.C.A.; supervision, H.C.L, L.K.M., and D.C.A.; funding acquisition, H.C.L., E.A.A, S.R., and D.C.A.

## ACKNOWLEDGEMENTS

We thank members of the Avgousti lab, M. Emerman, A. Geballe, M. Lagunoff, K. Lynch, S. Parkhurst, and M. Weitzman for insightful comments. We thank D. Janssens and the Henikoff lab for reagents and assistance with CUT&Tag. We thank the Salama and Overbaugh labs for assistance with ddPCR. We thank the Jerome lab for the clinical HSV isolates and the Cellular Imaging, Electron Microscopy (EMSR), F. Wu and Genomics & Bioinformatics Shared Resource Facilities at the Fred Hutchinson Cancer Research Center for help with sequencing and data analysis. We thank J. Dubrulle and the Cellular Imaging Shared Resource (CISR) for help with image analysis. The EMSR, CISR, and Genomics core are supported in part by the Fred Hutch/University of Washington Cancer Consortium (P30 CA015704). This study was supported by start-up funds from the RNA Bioscience Initiative at the University of Colorado School of Medicine (S. R.), the Fred Hutch (D.C.A.), and NIH funding to H.C.L. (AI083203), E.A.A. (AI083203), S.R. (GM133434), and D.C.A. (GM133441).

## MATERIALS AND METHODS

### Cells and viruses

Primary human foreskin fibroblasts (HFFs), hTERT-immortalized HFFs, and hTERT-immortalized macroH2A1 knockout HFFs were cultured using standard methods with 10% FBS and 1% penicillin-streptomycin as previously described^37^. Cells were grown at 37 °C with 5% CO_2_ and tested for mycoplasma contamination approximately once a month.

The lab adapted strain of HSV-1, *syn 17+* ^38^ was used for all experiments unless otherwise noted. Monolayers of cells were infected for 1 hour at 37 °C as previously described ^39^. An MOI of 3 was used for all experiments, except for EM samples which used MOI of 10. Cells were collected at mock, 4-, 8-, and 12-hpi for western blot. Supernatant was collected at 8- and 12-hpi for plaque assays. Virus stock was grown by infecting Vero cells at an MOI of 0.0001. Virus was harvested ∼60-80 hours post-infection and titered on U2OS cells to determine stock plaque forming units per mL (PFU/mL). Experimental plaque assays were set up in Vero cells. Plaque assays were set up on serial 10-fold dilutions in serum-free DMEM. Virus was left on cells for 1 hour then aspirated. Cells were washed with 1X PBS (pH 7.46) and 2% methylcellulose overlay in DMEM with 2% FBS and 0.5% penicillin-streptomycin was added to wells. Plaques were fixed with 0.2% crystal violet between 96-100 hours post infection and plaques were counted by hand. All plaque assays were set up with 2 technical replicates.

Clinical isolates were acquired in 1994 and 1995 from oral swab collections. Samples were de-identified on collection. Patients provided daily home collections of oral swabs, and qPCR was used to detect how many days HSV was detected in these swabs. Patients that had detectable HSV in only a few swabs were classified as “low-shedders” and patients that had a high percentage of days with positive swabs were classified as “high-shedders.” Stocks were a gift from the lab of Keith Jerome.

### Knockout of macroH2A1

MacroH2A1 guide RNA (gRNA, see Table 3) was cloned into TLCv2 (Addgene plasmid: 87360), a plasmid encoding doxycycline-inducible Cas9-2A-GFP and gRNA expression. To generate lentiviral particles, 1.0×10^7^ HEK293T cells were transfected with 12 µg TLCV2_sg_mH2A1, 8 µg pMDL (Addgene plasmid: 12251), 4 µg VSVg, and 2.5 µg pREV (Addgene plasmid: 115989) using Attractene transfection reagent (Qiagen: 301005). Lentivirus was harvested at 24-, 48-, and 72-hours post-transfection, filtered through 0.2 µm filters, aliquoted, flash frozen with liquid nitrogen, and stored at -80C. 2 mL of thawed filtered lentivirus was used to transduce approximately 2.25×10^6^ HFF-T or 3×10^6^ RPE cells in 10 cm plates with 8 µg/mL polybrene (Millipore-Sigma: TR-1003-G). Cells were allowed to reach confluence before selection with 1 µg/mL puromycin for 3 days. Cells were then sorted by GFP positivity or counted by hemocytometer, plated at 1 cell per well into 96-well plates, and selected through serial expansion of colonies. Selected cells were screened for macroH2A1 knockout by western blot.

**Table 1.**
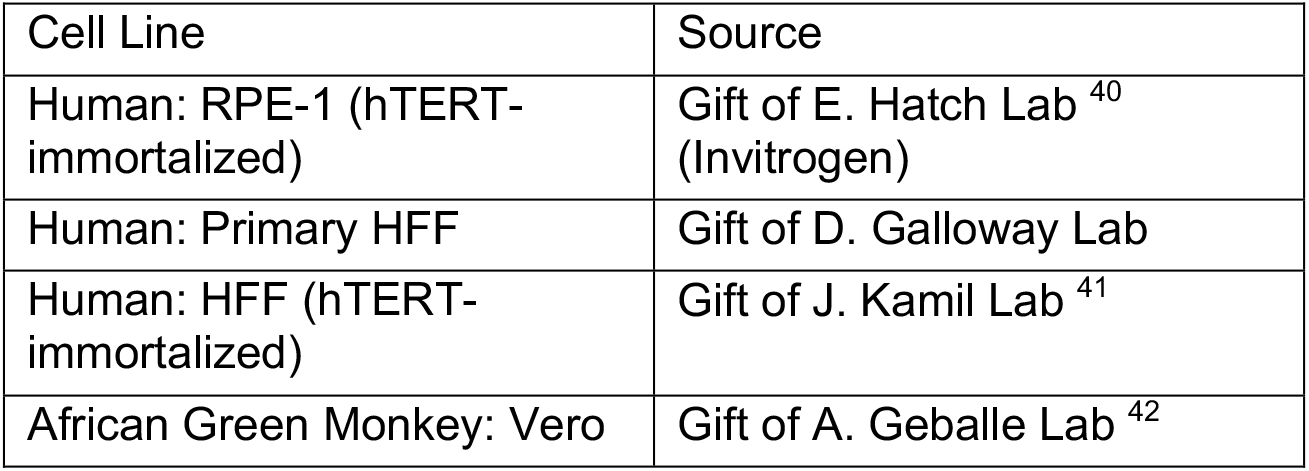
Cell line sources.

**Table 2.** Sources and identification of antibodies used.

| Antibody | Source | Identifier | Use (Concentration) |
| --- | --- | --- | --- |
| Mouse anti-actin | Abcam | Cat: 5441 | WB (1:10,000) |
| Mouse anti-gH | Abcam | Cat: 110227 | WB (1:1,000) |
| Rabbit anti-H3 | Abcam | Cat: 1791 | WB (1:20,000) |
| Rabbit anti-H3K27me3 | Cell Signal Technologies | Cat: 9733T | WB/CUT&Tag (1:100) |
| Rabbit anti-H3K9me3 | Abcam | Cat: 8898 | WB (1:1,000) |
| Rabbit anti-H4 | Abcam | Cat: 10158 | WB (1:1,000) |
| Mouse anti-ICP0 | Santa Cruz Biotechnology | Cat: 53070 | WB (1:1,000) |
| Rabbit anti-macroH2A1 | Abcam | Cat: 37264 | WB (1:1,000)<br>CUT&Tag (1:100) |
| Rabbit anti-macroH2A1.1 | Cell Signaling Technology | Cat: 12455S | WB (1:1,000) |
| Mouse anti-macroH2A1.2 | Cell Signaling Technology | Cat: 4827S | WB (1:1,000) |
| Rabbit anti-macroH2A2 | Active Motif | Cat: 39874 | WB (1:1,000) |
| HRP-conjugated anti-VP16 | Santa Cruz Biotechnology | Cat: 7546 | WB (1:1,000) |
| Peroxidase-AffiniPure Goat Anti-Mouse | Jackson Immunoresearch Laboratories | Cat: 115-035-003 | WB (1:5,000) |
| Peroxidase-AffiniPure Goat Anti-Rabbit | Jackson Immunoresearch Laboratories | Cat: 111-035-045 | WB (1:5,000) |
| Rabbit anti-mouse IgG | Abcam | Cat: 46540 | CUT&Tag (1:100) |
| Guinea pig anti-rabbit IgG | Antibodies Online | Cat: AB1N101961 | CUT&Tag (1:100) |

**Table 3.**
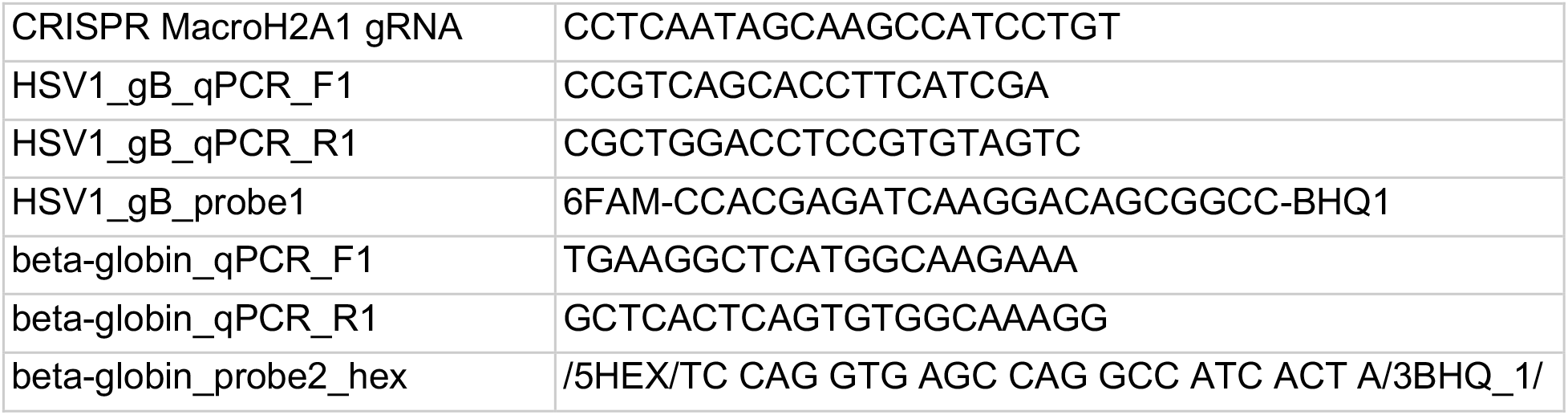
Primers used for knockout and ddPCR

### Infections with tazemetostat pretreatment

HFFs and macroH2A1 KO HFFs were treated with DMSO or 10 µM of tazemetostat (MedChem: HY-13803) in DMSO for 3 days prior to infection. Cells were then infected at MOI of 3 and after 1 hour of incubation with virus, fresh media with 10µM Taz was added to previously treated cells. Control samples were treated with equivalent volumes of DMSO. Samples were harvested as above.

### Western Blotting

Western blotting was performed as previously described ^37^. Briefly, cells were counted, pelleted, resuspended in 1x NuPAGE lithium dodecyl sulfate (LDS) sample buffer (Fisher Scientific: NP007) + 5% 2-betamercaptoethanol, and boiled for 15 minutes. Protein lysates were separated by 13.5% SDS-PAGE gels using NuPAGE MOPS buffer (Fisher Scientific: NP0001) at 75 volts for 30 minutes then 110 volts for 100 minutes, then transferred to a nitrocellulose membrane (Bio-Rad) at 100 volts for 70 minutes using Transfer Buffer (25 mM Tris Base, 100 mM glycine, 20% methanol). Membranes were ponceau stained and imaged. Membranes were blocked in 5% milk in Tris-buffered saline with Tween (TBST) for 1 hour, then probed with primary antibody overnight (see Table 2). Membranes were washed with TBST for thirty minutes, incubated with secondary antibodies conjugated to horseradish peroxidase (α-mouse or α-rabbit; 1:5000) at room temperature for 1 hour, washed with TBST for thirty minutes, and detected using Clarity Western ECL Substrate (Bio-Rad: 1705061) and Chemidoc MP Imaging System (Bio-Rad). Images were formatted using Adobe Photoshop and Illustrator.

### Quantification of HSV-1 genomes by ddPCR

Cells were harvested at the indicated times post-infection by trypsinization, washed with 1X PBS and centrifuged at 5000 xg for 2 minutes. Pellets were flash-frozen in liquid nitrogen and stored at -80 °C until processed. HSV-1 DNA within cells was isolated from frozen pellets using QIAamp DNAMini Kit (Qiagen: 51304).

Supernatants were harvested at the indicated times post-infection, centrifuged at > 3500 xg and filtered through 40 micron sterile syringe filters. DNA on the exterior of filtered capsids was digested for 1 hour at 25 °C with 20.3 units DNase (Qiagen: 79254) supplemented with 10 mM MgCl_2_. DNase was inactivated at 75 °C for 10 minutes followed by vortexing. Capsids were digested with 3 mg/mL proteinase K (Fisher Scientific: BP1700) in 100 mM KCl, 25 mM EDTA, 10 mM Tris-HCl pH 7.4 and 1% Igepal for 1 hour at 50 °C. HSV-1 genomes from digested capsids were isolated using QIAamp DNAMini Kit.

A duplexed droplet digital PCR was performed to measure the levels of cellular or supernatant HSV-1 genomes on the QX100 droplet digital PCR system (Bio-Rad Laboratories, Hercules, CA) using a primer/probe set specific to HSV-1 gB. Cell numbers were determined using a primer/probe set specific to human Beta-globin, a reference gene that exists at two copies per cell. See Table 3 for oligonucleotide sequences, as have been previously established for HSV-1_43_. The digital droplet PCR (ddPCR) reaction mixture consisted of 12.5 µL of a 2X ddPCR Supermix for Probes no dUTP (Bio-Rad: 1863024), 1.25 µL of each 20X primer-probe mix, and 10 µL of template DNA. 20 µL of each reaction mixture was loaded onto a disposable plastic cartridge (Bio-Rad: 1864008) with 70 µL of droplet generation oil (Bio-Rad: 1863005) and placed in the droplet generator (Bio-Rad). Droplets generated were transferred to a 96-well PCR plate (Bio-Rad 12001925), and PCR amplification was performed on a Bio-Rad C1000 Touch Thermal Cycler with the following conditions: 95 °C for 10 minutes, 40 cycles of 94 °C for 30 seconds, and 60 °C for 1 minute, followed by 98 °C for 10 minutes and ending at 4 °C. After amplification, the plate was loaded onto the droplet reader (Bio-Rad: QX200), and the droplets from each well of the plate were automatically read with droplet reader oil (Bio-Rad: 186-3004) at a rate of 32 wells per hour. Data were analyzed with QuantaSoft analysis software and quantitation of target molecules presented as copies per µL of PCR reaction. HSV-1 genome values were standardized to cellular β-globin levels. Experiments were completed in biological triplicate and statistical analysis was performed using Prism v10 (GraphPad Software).

### Cleavage Under Targets & Tagmentation (CUT&Tag)

Two biological replicates per time point were obtained from independent infections. Protocol was adapted from the established CUT&Tag methods^20^. Cells were harvested using trypsin, washed three times with ice-cold phosphate-buffered saline (PBS) via centrifugation at 600 xg for 3 minutes and counted using a hemocytometer. Nuclei from 600,000 cells were isolated by hypotonic lysis in 1 mL buffer NE1 (20 mM HEPES-KOH pH 7.9; 10 mM KCl; 0.5 mM spermidine; 0.1% Triton X-100; 20% Glycerol; Roche EDTA-free protease inhibitor) for 10 minutes on ice followed by centrifugation at 1300 xg for 4 minutes. Nuclei were resuspended in Wash buffer (20 mM HEPES-NaOH pH 7.5; 150 mM NaCl; 0.5 mM spermidine; Roche EDTA-free protease inhibitor) and counted using a hemocytometer. BioMag Plus Concanavalin A coated beads (Polysciences: 86057-3) were equilibrated with Binding buffer (20 mM HEPES-KOH pH 7.9; 10 mM KCL; 1mM CaCl_2_; 1 mM MnCl_2_). Beads (5 µL) were mixed with aliquots of 75,000 nuclei and incubated at 25 °C for 10 minutes followed by magnetic separation of beads. Beads were resuspended in 50 µL primary antibody [anti-mH2A1 (Abcam: ab37624), anti-H3K27me3 (Cell Signaling:9733) or anti-mouse IgG (Abcam: ab46540) in Wash buffer supplemented with 2 mM EDTA and 0.1% bovine serum albumin (BSA) and incubated on a nutator at 4 °C overnight. The beads were decanted on a magnet stand then resuspended in 50 µL secondary antibody [Guinea pig anti-rabbit IgG (Antibodies-Online: ABIN101961) 1:100] in Wash buffer supplemented with 2 mM EDTA and 0.1% BSA and incubated on a nutator at room temperature for 1 hour. The beads were decanted on a magnet stand and washed with 200 µL Wash buffer, then were resuspended in 50 µL pA-Tn5 (1:200) in 300-Wash buffer (Wash buffer containing 300 mM NaCl) and incubated on a nutator at room temperature for 1 hour. The beads were washed twice with 200 µL 300-Wash buffer, then resuspended in 50 µL Tagmentation buffer (300-Wash buffer supplemented with 10 mM MgCl_2_) and incubated at 37 °C for 1 hour. Beads were then washed with 50 µL TAPS wash buffer (10 mM TAPS pH 8.5, 0.2 mM EDTA), then resuspended in 5 µL TAPS wash buffer supplemented with 0.1% SDS and incubated at 58 °C for 1 hour. SDS was neutralized on ice with 15 µL 0.67% Triton-X100. 2 µL of 10 mM indexed P5 and P7 primer solutions and 25 µL NEBnext High-Fidelity 2X Master Mix (New England BioLabs: ME541L) were added. Gap-filling and 15 cycles of PCR were performed using an MJ PTC-200 Thermocycler. Library clean-up was performed by incubating beads with 65 µL SPRI bead slurry for 5-10 minutes, then magnetization and two washes with 200 µL 80% ethanol. Libraries were eluted with 22 µL Tris-HCl pH 8.0 and 2 µL was used for Agilent 4200 Tapestation analysis. The barcoded libraries were mixed to achieve equimolar representation as desired aiming for a final concentration as recommended by the manufacturer for sequencing on an Illumina HiSeq 2500 2-lane Turbo flow cell.

### CUT&Tag data processing

CUT&Tag raw sequencing data were aligned to a custom genome made by concatenating human (hg38), HSV (JN555585.1), and E. coli (U00096.3 Escherichia coli str. K-12 substr. MG1655, complete genome). We performed alignments using bowtie2.

### Domain calling

Coverage at 100 bp windows of the human hg38 reference genome was calculated as number of reads of given length (120-1000 bp for the analyses presented here) that mapped at that window normalized by the factor N:

N= 10,000/(Total number of reads that mapped to E. coli genome)

Here 10,000 is an arbitrarily chosen number. We used *E. coli* DNA as a spike-in to normalize all datasets. We partitioned all chromosomes into domains of macroH2A1: domains had an enrichment that was two times the genome-wide median and at least four-fold higher than the IgG control. The normalized coverage at each base-pair from each replicate was averaged when combining multiple replicates. The normalized read density in 100 bp bins were then smoothed with a running average over the bins spanning +/-1000 bp around each bin. We then calculated the genome-wide distribution of normalized read density and medians that were plotted in Figure S1. We averaged medians across WT macroH2A1 datasets (mock, 4, 8, and 12hpi), and multiplied the average by 2, to set the cutoff for domain definition for macroH2A1 datasets. We defined a similar cutoff for H3K27me3 datasets using H3K27me3 WT datasets.

### Identifying domain level dynamics of macroH2A1 over time course of infection

To measure changes in macroH2A1 across time points where the domain boundaries are not the same, we first concatenated domain definitions from all macroH2A1 datasets and then defined a set of non-overlapping intervals using the “disjoin” method of GenomicRanges R package ^44^. We then calculated the log2 ratio of macroH2A1 enrichment over IgG for the non-overlapping regions for the mock, 4, 8, and 12hpi. The 4, 8, and 12hpi enrichments were then divided by the enrichment of the mock dataset to obtain change in macroH2A1 over time course at the non-overlapping regions. k-means clustering (k=6) using R ^45^ was performed on the matrix where the rows are the non-overlapping regions and the columns are change in macroH2A1 over mock at 4, 8, and 12hpi. We extended the non-overlapping regions by 5 bp on each end and then merged regions within each cluster using “reduce” method of GenomicRanges, to obtain domains in each cluster. We recalculated change in macroH2A1 at 4, 8, and 12hpi over mock, which was used to plot heatmaps and boxplots shown in Figure 1 and Figure S1. H3K27me3 enrichment for WT and macroH2A1 KO cells was calculated at clustered macroH2A1 domains. The unique code for this work can be found here: https://doi.org/10.5281/zenodo.6783949

### RNA sequencing

Three biological replicates per time point were obtained from independent infections. Cells were harvested at the indicated times post-infection by trypsinization, washed with PBS and centrifuged at 5000 xg for 2 minutes. Pellets were lysed with TRIzol (Thermo Fisher: 15-596-026) and total RNA was harvested according to manufacturer’s instructions. RNAs were then treated with DNase (Qiagen: 79254) on RNeasy columns (Qiagen: 74104) per manufacturer’s instructions. RNA was precipitated with 3 volumes ice-cold 96% ethanol, 1 volume 3 M sodium acetate pH 5.5, and 1 µL glycogen (Thermo Fischer: R055) overnight at -80 °C. Precipitated RNAs were pelleted at 15,000 xg and 4 °C for 30 minutes, washed with ice-cold 75% ethanol and spun as above for 10 min. RNA was resuspended in nuclease-free water.

RNA was quantified by Nanodrop and integrity analyzed with the 4200 Tapestation Bioanalyzer system (Agilent). 500 ng of total RNA with an RNA Integrity Number (RIN) greater than 9.5 were used to prepare sequencing libraries with the TruSeq® Stranded mRNA Library Prep Kit (Illumina: 20020594). Library concentrations were measured with Qubit dsDNA HS Assay Kit (Thermo Fisher: Q32854) then analyzed with Agilent High Sensitivity D5000 ScreenTape System and pooled. Libraries were sequenced with 100-bp paired-end reads on an Illumina NovaSeq 6000 SP sequencer at the Fred Hutch Genomics Core Facility

### RNA/4sU-RNA/Hi-C data processing

Fastq files were filtered to exclude reads that didn’t pass Illumina’s base call quality threshold. STAR v2.7.1 ^46^ with 2-pass mapping was used to align paired-end reads to a combined reference of human genome build hg38 and HSV1 genome JN555585.1 (https://www.ncbi.nlm.nih.gov/nuccore/JN555585.1/). FastQC 0.11.9 (https://www.bioinformatics.babraham.ac.uk/projects/fastqc/) and RSeQC 4.0.0 ^47^ were used for QC including insert fragment size, read quality, read duplication rates, gene body coverage and read distribution in different genomic regions. FeatureCounts ^48^ in Subread 1.6.5 was used to quantify gene-level expression by strand-specific paired-end read counting.

Gene annotation were based on GENCODE V31 (https://www.gencodegenes.org/human/) and GCA_000859985.2_ViralProj15217 (https://www.ncbi.nlm.nih.gov/data-hub/genome/GCF_000859985.2/). For HSV1, annotated genes were collapsed into non-overlapping transcribed regions, e.g. X indicating a transcribed region unique to gene X, X:Y indicating an overlapping transcribed region for genes X and Y, and so on.

Bioconductor package edgeR 3.26.8 ^49^ was used to detect differential gene expression between sample groups. Genes with low expression were excluded by requiring at least one count per million in at least N samples (N is equal to the number of samples in the smaller group). The filtered expression matrix was normalized by TMM method (https://genomebiology.biomedcentral.com/articles/10.1186/gb-2010-11-3-r25) and subject to significance testing using quasi-likelihood pipeline implemented in edgeR. Genes were deemed differentially expressed if fold changes were greater than 2 in either direction and Benjamini-Hochberg adjusted p-values were less than 0.01.

The intervals representing start and end of each gene were intersected with clustered macroH2A1 domains to obtain cluster assignments for genes. Intersection was performed using “intersect” function of bedtools ^50^. Genes that did not uniquely intersect with domains in a single cluster were discarded. The cluster assignments for genes were used for plotting total RNA fold changes in Figure 2. Reads per kilobase of transcript per million reads mapped (RPKM) values for 4sU-RNA was obtained from GEO (GSE59717) and converted to transcripts per million (TPM). The TPM values across two 4sU-RNA replicates for each condition was averaged and box plots were generated similarly to total RNA.

For calculating eigenvector distributions, compartment wig file was downloaded from the following link: https://4dn-open-data-public.s3.amazonaws.com/fourfront-webprod/wfoutput/b543cbf4-ce54-4d2d-8960-281528ff18a6/4DNFI342UZP1.bw

Regions from compartment file intersecting with each domain cluster were extracted and the values were used for generating box plots shown in Figure 2D.

### Electron Microscopy

Cells were fixed in 2% paraformaldehyde and 2.5% glutaraldehyde in 0.1M sodium cacodylate buffer (pH 7.3) at 4 °C. Fixed cells were rinsed briefly in 1% sucrose in 50mM cacodylate (pH 7.2), then postfixed on ice for 30 mins in a solution of 1% osmium (EM Sciences: RT19152) and 0.8% potassium ferricyanide in 50mM cacodylate (pH 7.2). Cell pellets were washed twice briefly at 25 °C in 1% sucrose in 50mM cacodylate (pH 7.2), then washed in three changes of 50mM cacodylate (pH 7.2) for 5 min each. Cell pellets were treated with 0.2% tannic acid (Sigma-Aldrich: 1401-55-4) in 50 mM cacodylate (pH 7.2) for 15 minutes at 25 °C, then rinsed several times in water. Cells were dehydrated through a graded ethanol series and embedded in Epon 12 resin (Ted Pella: 18010). 70nm thin sections were cut using an Ultracut UC7 ultramicrotome (Leica Mikrosysteme) and collected on 200 mesh formvar/carbon copper grids (Ted Pella: 01800). Sections were stained with 2% aqueous uranyl acetate and Reynolds lead citrate. Cell pellet sections were imaged using a Talos L120C microscope operated at 120kV with a Ceta-16M (4096 × 4096) camera (ThermoFisher Scientific).

All data were collected at spot size 5 with a 100um C2 aperture and 70um objective aperture. For quantification, nuclei were targeted at 1250x and manually circled. Autofocus was set to - 2.0um. Nuclei were then imaged at 11000x as a montage and stitched together automatically using SerialEM (Nexperion Inc). Stitched maps were exported as uncompressed 16-bit tif files for further analysis. For qualitative analysis, images were manually focused to -2.0um defocus and then 20 1-second frames were collected and drift-corrected using Velox software (Thermo Fisher Scientific, Eindhoven, NL). Final summed images were exported as 16-bit tif files and cropped using ImageJ.

Image analysis pipelines for counting capsids and for measuring chromatin density at the nuclear envelope were deployed in MATLAB R2020b. Scripts are available on the Fred Hutch GitHub repository. Capsid pipeline follows three steps: 1-nuclear boundaries identification, 2-capsid detection, 3-capsid classification. Nuclear boundaries were identified with user input, by outlining a freehand contour of the nucleus of interest, from which a binary mask is extracted. Capsid detection was performed on the median-filtered complement of the original image, using a Circular Hough Transform based algorithm, with phase coding for radii estimation^51^, and search radius ranging from 10.6 to 31.7 nm. Capsids residing within the nuclear mask were then counted and classified. Detected capsids were classified in three categories (Empty, Intermediate and Full), depending on the distribution of the pixel grayscale intensities within the capsids relative to a normal distribution.

Width of heterochromatin abutting the nuclear envelope was quantified by measuring the length of the binarized chromatin from 1D intensity profiles along the normal of the nuclear perimeter, sampled at every 10 perimeter pixels. Dense heterochromatin was binarized using global Otsu’s thresholding method applied to the background-corrected complement of the contrast-adjusted original image. Noise from the binarized image was further reduced by applying a 2D order statistic filter using the minimum value of a varying domain interactively defined by the user, with default value of 8-by-8 pixels. The resulting heterochromatin density distribution was normalized to the total length of the nucleus’ perimeter.

## SUPPLEMENTAL FIGURES

**Supplemental Figure 1.**
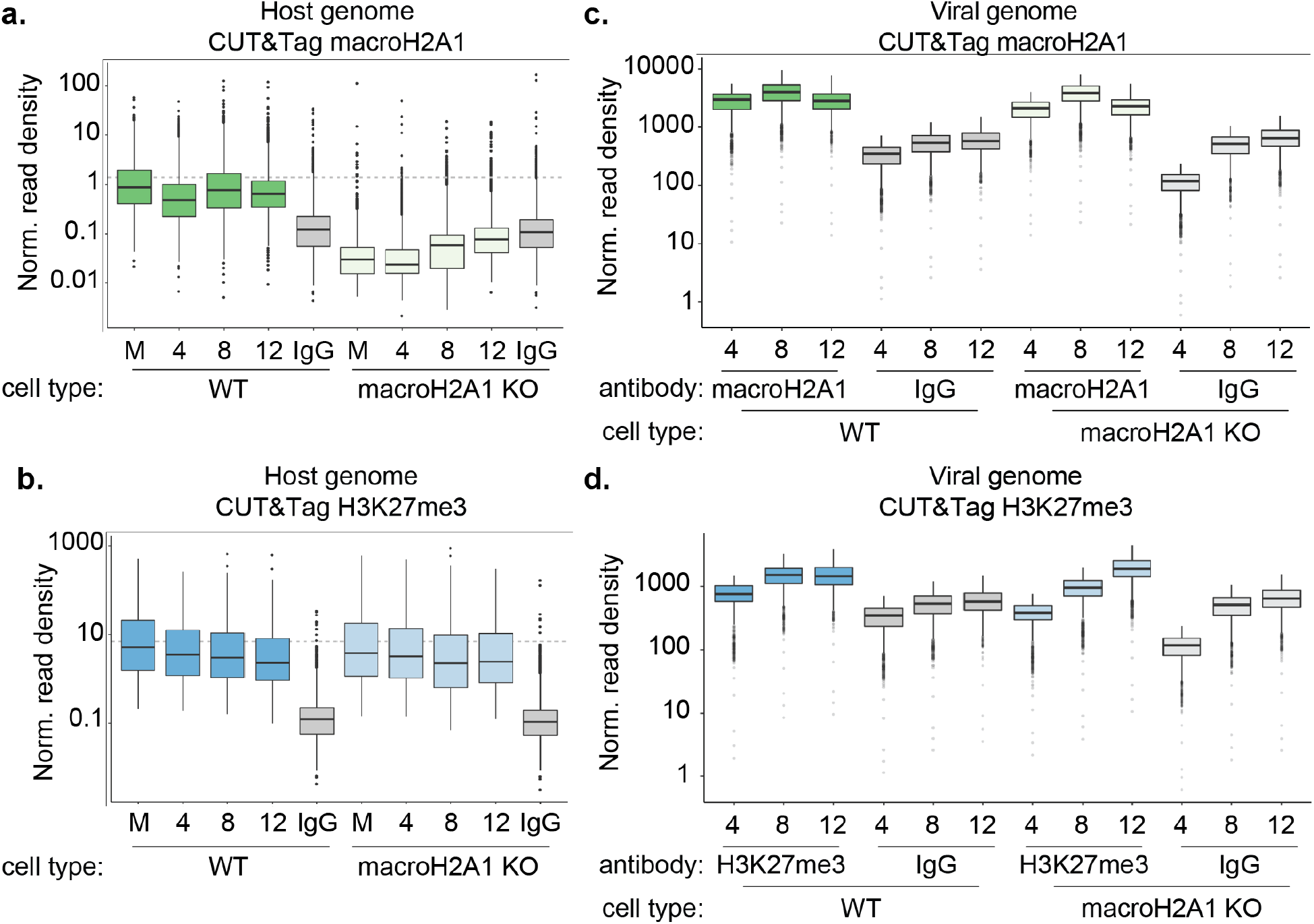
Quantification of macroH2A1 and H3K27me3 enrichment on host and viral genomes during HSV-1 infection. a) Spike-in normalized CUT&Tag read density of macroH2A1 on WT HFF-T and macroH2A1 KO cell genomes during HSV-1 infection. b) Spike-in normalized CUT&Tag read density of H3K27me3 on WT HFF-T and macroH2A1 KO cell genomes during HSV-1 infection. c) Spike-in normalized CUT&Tag read density of macroH2A1 on HSV-1 genomes during infection of WT HFF-T and macroH2A1 KO cells. d) Spike-in normalized CUT&Tag read density of H3K27me3 on HSV-1 genomes during infection of WT HFF-T and macroH2A1 KO cells.

**Supplemental Figure 2.**
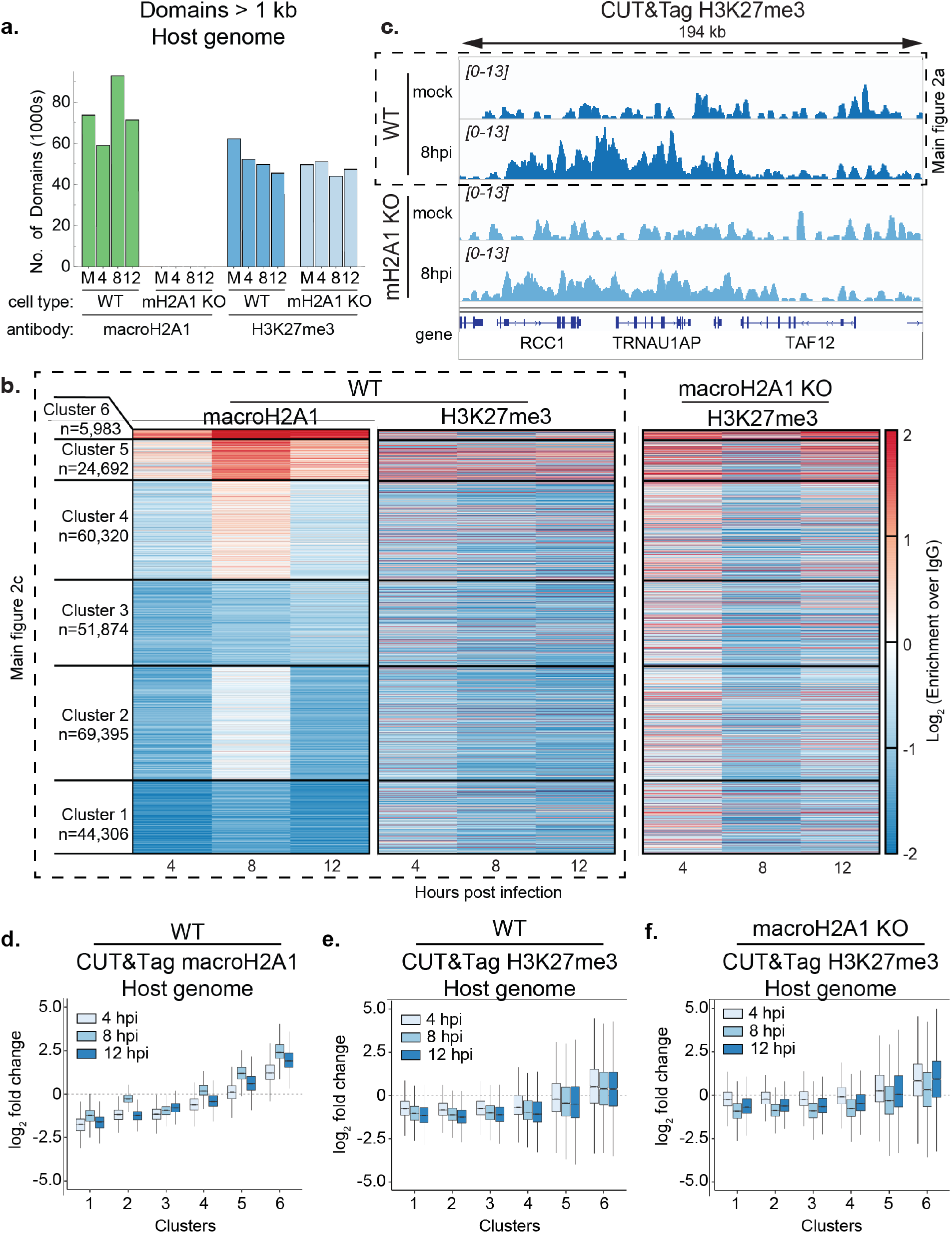
Quantification of macroH2A1 and H3K27me3 bound regions on host genomes during HSV-1 infection. a) Total number of macroH2A1 and H3K27me3 domains over 1 kb as measured by CUT&Tag of WT HFF-T and macroH2A1 KO cell genomes during HSV-1 infection. b) Heat maps of macroH2A1 and H3K27me3 dynamics over the course of HSV-1 infection generated from CUT&Tag data. c) Representative screenshots of H3K27me3 broad domain increase over the course of HSV-1 infection as measured by CUT&Tag. d) Quantification of heat maps from Figure S2B showing macroH2A1 enrichment across the clusters. e) Quantification of heat maps from Figure S2B showing H3K27me3 enrichment in clusters defined by macroH2A1 as in D. f) Quantification as in E in macroH2A1 KO cells.

**Supplemental Figure 3.**
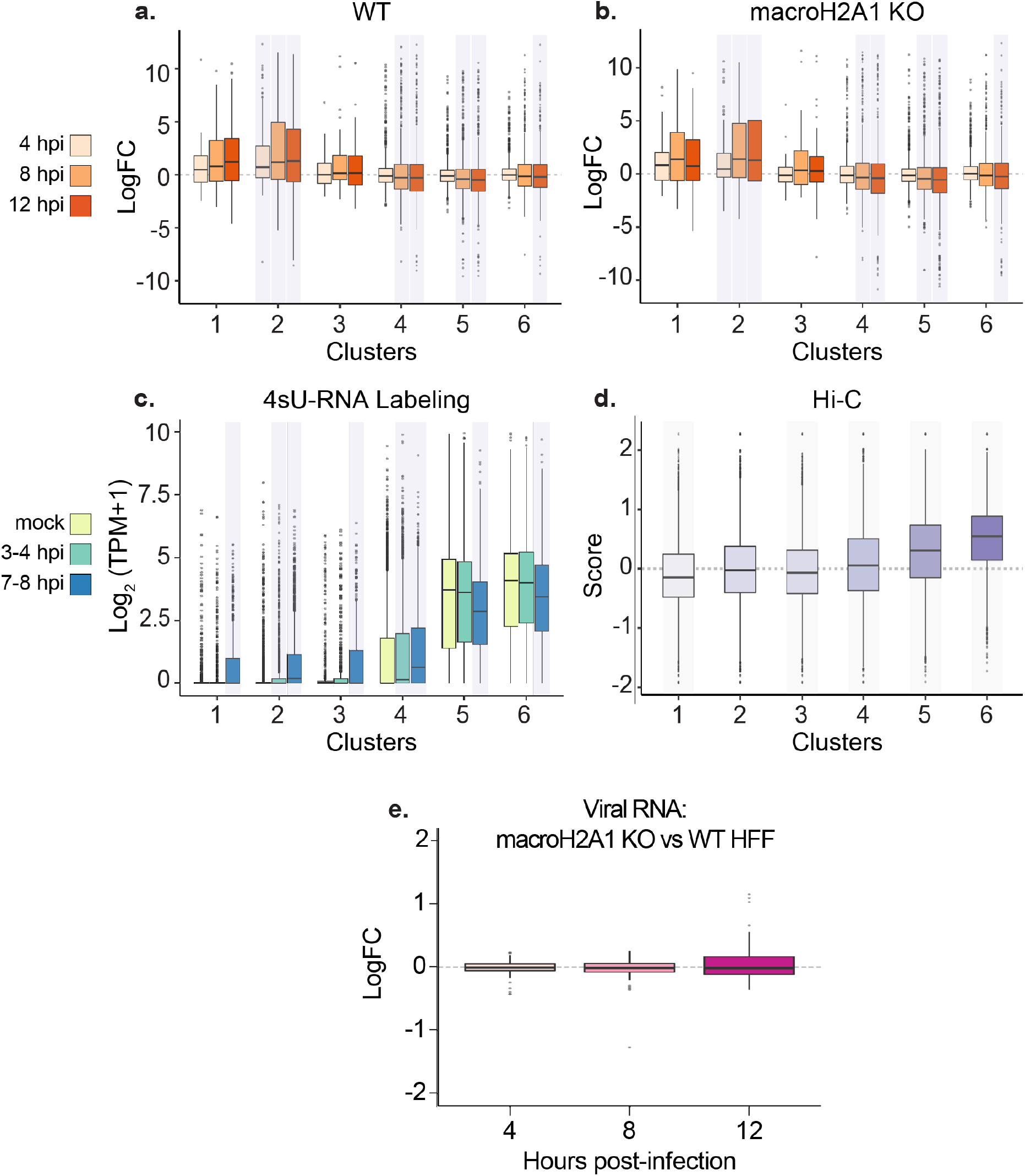
MacroH2A1 and H3K27me3 presence on host genomes correlates with decreased transcription. a) Box plots of total host RNA from RNA-seq in HFF-T WT for genes overlapping clusters from Figure 2. Kolmogrov-Smirnov test (in R) was performed comparing logCPM values of genes at each time point with logCPM values of genes from mock dataset for each cluster. The p-values were corrected for multiple testing, and the time points and clusters with corrected p-value less than 0.05 are shaded. b) Same as (a) for macroH2A1 KO cells. c) 4sU-RNA counts for genes from Hennig *et al*^30^ overlapping with clusters from Figure 2 shown as box plots and shading as indicated in (a). d) Box plots of Hi-C eigenvector scores of regions overlapping with clusters from Figure 2. The Hi-C compartment eigen vector scores were obtained from 4DN project website. Wilcoxon signed rank test with alternate hypothesis that the true location is not equal to 0 was performed on the distribution of eigen vector scores for each cluster. The p-values were corrected for multiple testing, and the clusters with corrected p-value less than 0.05 are shaded. Bonferroni correction was used for p-value adjustments. e) Log2 fold-change of HSV-1 transcripts at 4, 8 and 12 hpi identified by RNA-seq in macroH2A1 KO compared to WT HFF-T cells. There is no significant difference between the cell types.

**Supplemental Figure 4.**
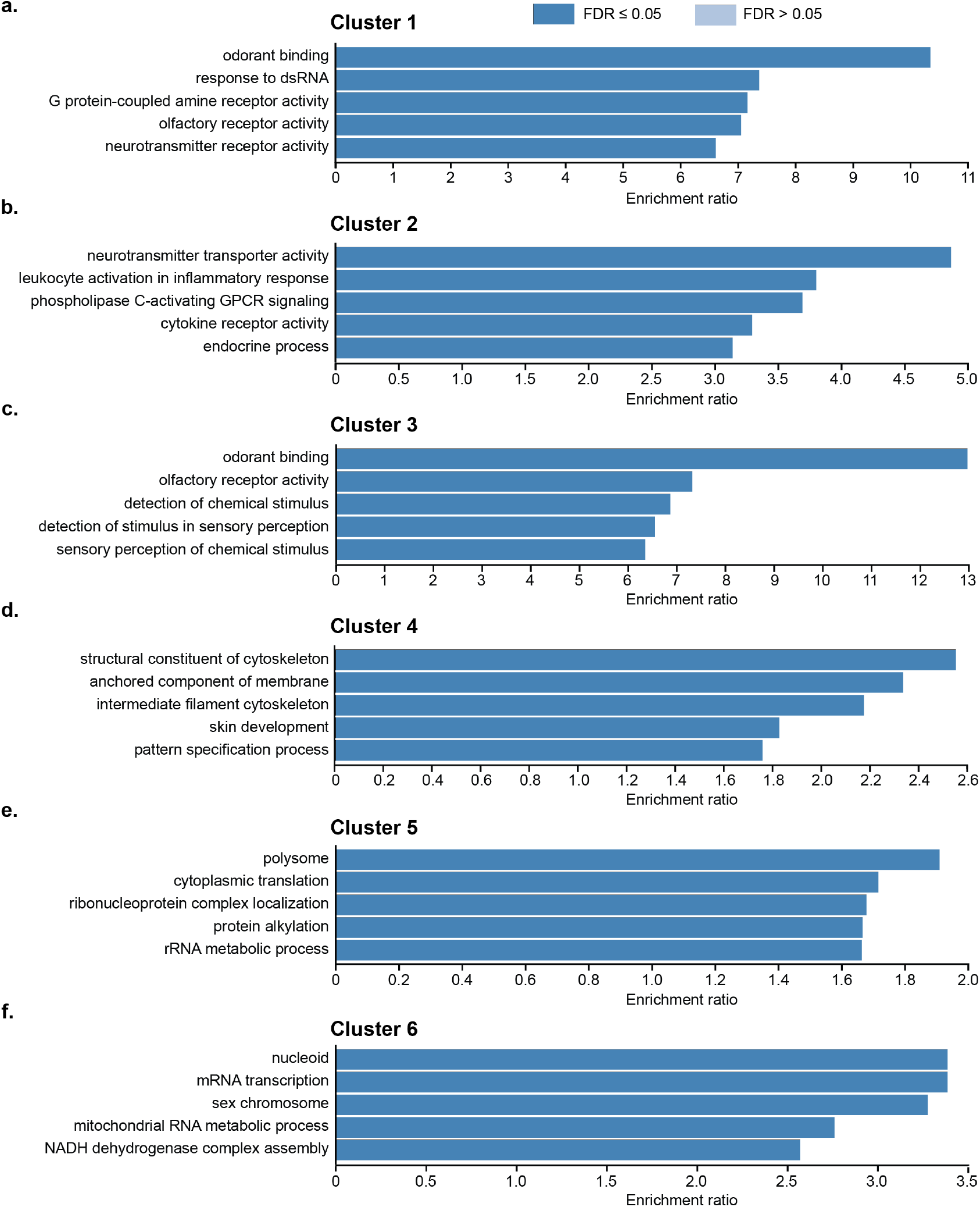
Gene ontology enrichment analysis of total host mRNA defined by macroH2A1 clusters. Gene ontology analysis of mRNA levels between WT HFFs and macroH2A1 KO cells. Cluster 1 (a), cluster 2 (b), and cluster 3 (c), cluster 4 (d), cluster 5 (e), and cluster 6 (f) were defined by macroH2A1 domains in Figure 1. P-value <0.001 for all shown GO clusters and FDR was <0.05.

**Supplemental Figure 5.**
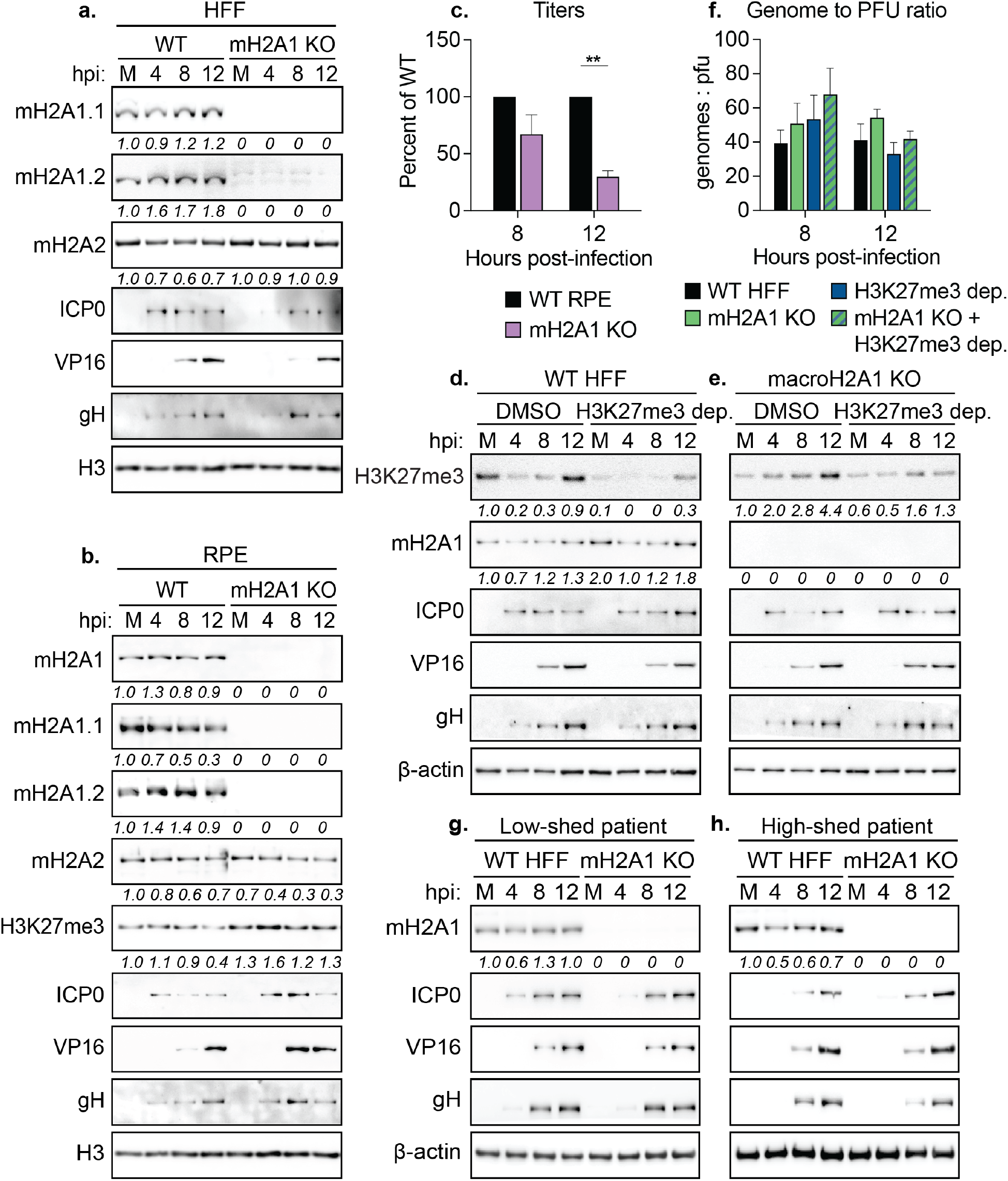
Viral protein levels do not change upon macroH2A1 knockout or H3K27me3 depletion in HFFs and RPEs. a) Western blot of total macroH2A1, macroH2A1.1, and macroH2A1.2 isoforms and macroH2A2 in mock(M) or during HSV-1 infection at 4-, 8-, and 12-hpi in WT and macroH2A1 KO HFF-T cells. H3 is shown as a loading control. Quantification of relative intensity over H3 normalized to the mock samples are shown. b) Western blot as in (a) in WT and macroH2A1 KO tert-immortalized RPE cells. H3 is shown as a loading control. c) Infectious progeny produced by WT or macroH2A1 knockout RPE cells quantified by plaque assay. Viral yield calculated as in Fig 3c for conditions as indicated, *** p. < 0.001. d) Western blot of H3K27me3 and macroH2A1 during HSV-1 infection shown as mock infected (M), 4-, 8-, and 12-hpi. WT cells H3K27me3 depleted or treated with DMSO for 3 days prior to infection and remained with inhibitor over the course of infection. Beta-actin is shown as a loading control. Quantification of relative intensity over beta-actin normalized to the mock sample on the same blot are shown. e) Western blot as in (d) for macroH2A1 KO cells. f) Western blots as in (a) for infections with low-shedding clinical isolate of HSV-1. Fold-change of macroH2A1 protein levels normalized to macroH2A1 level in mock-infected cells. g) Western blot as in (f) for high-shedding clinical HSV-1 isolate.

**Supplemental Figure 6.**
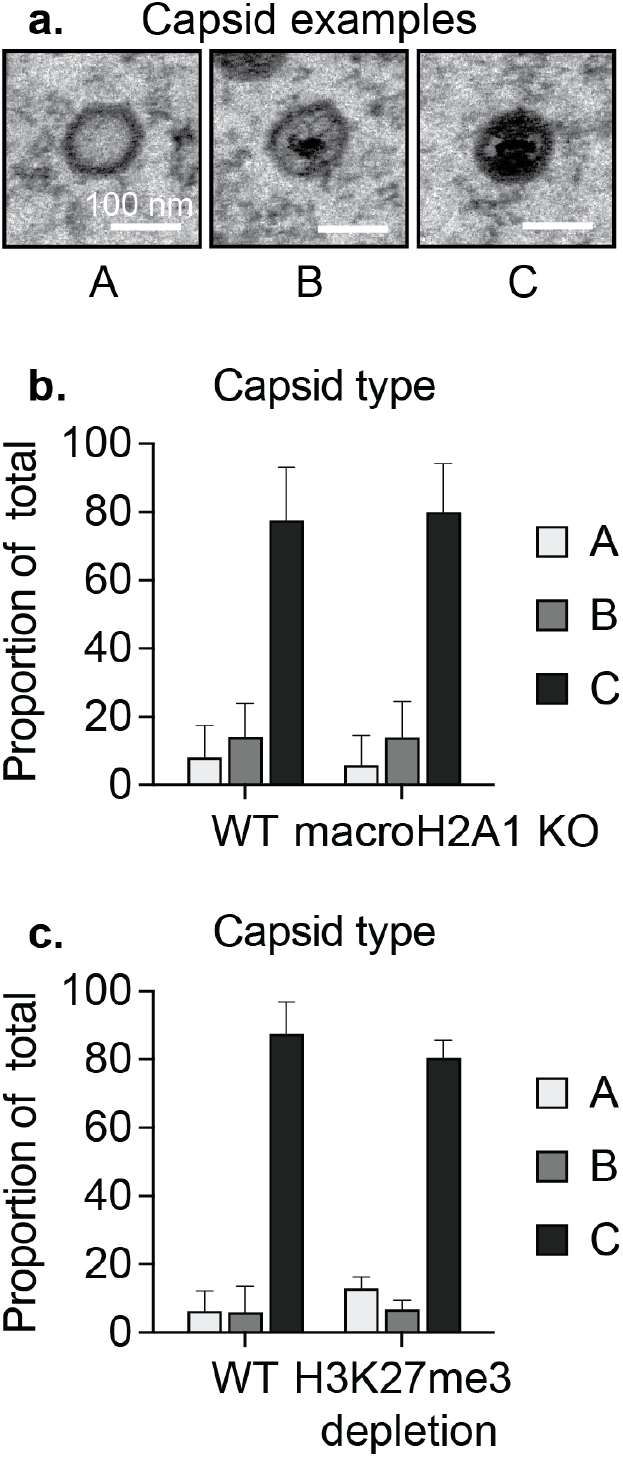
HSV-1 capsid formation is independent of macroH2A1. a) TEM images of representative A (empty), B (scaffolding proteins, but no genome), and C (full) HSV-1 capsids. Scale bars as noted for all TEM images. b) Quantification of capsid type within nuclei from (a) in WT and macroH2A1 KO cells. Values for each capsid type are shown as a proportion of total capsids. No significance by ANOVA. c) Capsid type quantification as in (b) for H3K27me3 depleted cells.

